# Chloroplast-derived photo-oxidative stress causes changes in H_2_O_2_ and *E*_GSH_ in other subcellular compartments

**DOI:** 10.1101/2020.07.20.212670

**Authors:** José Manuel Ugalde, Philippe Fuchs, Thomas Nietzel, Edoardo A. Cutolo, Ute C. Vothknecht, Loreto Holuigue, Markus Schwarzländer, Stefanie J. Müller-Schüssele, Andreas J. Meyer

## Abstract

Metabolic fluctuations in chloroplasts and mitochondria can trigger retrograde signals to modify nuclear gene expression. Mobile signals likely to be involved are reactive oxygen species (ROS), which can operate protein redox switches by oxidation of specific cysteine residues. Redox buffers such as the highly reduced glutathione pool serve as reservoirs of reducing power for several ROS scavenging and ROS-induced damage repair pathways. Formation of glutathione disulfide (GSSG) and a shift of the glutathione redox potential (*E*_GSH_) towards less negative values is considered a hallmark of several stress conditions. Here we used the herbicide methyl viologen (MV) to generate ROS locally in chloroplasts of intact Arabidopsis seedlings and recorded dynamic changes in *E*_GSH_ and H_2_O_2_ levels with the genetically-encoded biosensors Grx1-roGFP2 (for *E*_GSH_) and roGFP2-Orp1 (for H_2_O_2_) targeted to chloroplasts, the cytosol or mitochondria. Treatment of seedlings with MV caused a rapid oxidation in chloroplasts and subsequently also in the cytosol and mitochondria. The MV-induced oxidation was significantly boosted by illumination with actinic light and largely abolished by inhibitors of photosynthetic electron transport. In addition, MV also induced an autonomous oxidation in the mitochondrial matrix in an electron transport chain activity-dependent manner that was milder than the oxidation triggered in chloroplasts by the combination of MV and light. *In vivo* redox biosensing resolves the spatiotemporal dynamics of compartmental responses to local ROS generation and provide a basis for understanding how compartment-specific redox dynamics may operate in retrograde signaling and stress acclimation in plants.

**One sentence summary:** Methyl viologen-induced photooxidative stress causes an increase of H_2_O_2_ and oxidation of glutathione in chloroplasts, cytosol and mitochondria as well as autonomous oxidation in mitochondria.

## INTRODUCTION

Communication between different subcellular compartments of plant cells is fundamental to establish and sustain cooperative functioning and to acclimate to diverse environmental conditions. Since most plastidial and mitochondrial proteins are encoded in the nuclear genome, retrograde signals from the organelles to the nucleus are essential to adjust organelle function by coordinating the expression of nuclear and organellar genomes (Van Aken et al., 2016; de Souza et al., 2017; Dietz et al., 2019). Communication between the endosymbiotic organelles and the nucleus is likely to involve the cytosol as the intermediate compartment. However, chloroplasts can also make direct physical contact with the nuclear envelope via stromules, which has been suggested to mediate signaling (Caplan et al., 2015; Erickson et al., 2017; Exposito-Rodriguez et al., 2017). Physical interaction also occurs between different organelles and may facilitate efficient exchange of metabolites and information (Pérez-Sancho et al., 2016; Perico and Sparkes, 2018).

Reactive oxygen species (ROS), such as H_2_O_2_, have emerged as signaling molecules in plants and their roles in early signaling events initiated by cellular metabolic perturbation and environmental stimuli are established (Waszczak et al., 2018; Smirnoff and Arnaud, 2019). During unfavorable environmental conditions, superoxide (O_2_^·-^) is produced at an increased rate by the electron transport chains (ETCs) in chloroplasts and mitochondria. O_2_^·-^ is rapidly converted to hydrogen peroxide (H_2_O_2_) and molecular oxygen (O_2_) by superoxide dismutases (SODs). H_2_O_2_ can be further detoxified through a set of peroxidases, including peroxiredoxins (PRX) (Liebthal et al., 2018)) several glutathione *S*-transferases (GST) (Sylvestre-Gonon et al., 2019; Ugalde et al., 2020), glutathione peroxidase-like enzymes (GPXL) (Attacha et al., 2017), and ascorbate peroxidases (APX). The latter operate as part of the glutathione–ascorbate cycle in the plastid stroma, mitochondrial matrix, peroxisomes and cytosol (Foyer and Noctor, 2005; Narendra et al., 2006). The transient drain of electrons from the local glutathione redox buffer causes a concomitant increase in glutathione disulfide (GSSG) and hence a change in the glutathione redox potential *E*_GSH_) (Marty et al., 2009; Bangash et al., 2019; Nietzel et al., 2019; Wagner et al., 2019). Intracellular H_2_O_2_ levels reached under stress conditions can affect cellular redox regulation leading to the oxidation of protein thiols (Dietz et al., 2016). H_2_O_2_ was shown to diffuse across the chloroplast envelope even at low concentrations, and it has been estimated that about 5% of the total ROS produced in high light leave the chloroplast (Mubarakshina et al., 2010). Those properties contributed to the suggestion of H_2_O_2_ to operate as a messenger in signaling processes arising from the organelles. Moreover, direct transfer of H_2_O_2_ from a subpopulation of chloroplasts localized in close proximity to the nucleus itself was recently found to mediate photosynthetic control over gene expression in tobacco leaves (Caplan et al., 2015; Exposito-Rodriguez et al., 2017).

ROS production at specific sites of the photosynthetic ETC (pETC) can be artificially enhanced by using inhibitors and redox catalysts. Among these, the herbicide methyl viologen (MV) acts by re-directing electrons from photosystem I (PSI) to O_2_ and thereby enhancing the production of O_2_^·-^ (Scarpeci et al., 2008). Based on its mechanism, MV is also useful as an experimental cue to induce photo-oxidative stress in photosynthetic organisms. In mammals and other non-photosynthetic organisms, MV induces the generation of O_2_^·-^ by re-directing electrons from complex I of the mitochondrial ETC (mETC) to O_2_ (Cochemé and Murphy, 2008), suggesting that current models to study retrograde signaling are likely to be more complex than previously expected and involve additional subcellular sites (Cui et al., 2019; Shapiguzov et al., 2019). Steady-state measurements in cotyledons of Arabidopsis seedlings have previously shown that MV can induce oxidation in both cytosol and mitochondria in the absence of illumination (Schwarzländer et al., 2009).

Chemical probes for detecting ROS in living systems, such as 2’,7’-dihydrodichlorofluorescein diacetate are typically converted to a fluorescent product through reaction with ROS and accumulate in tissues with different specificities for distinct forms of ROS (Fichman et al., 2019). While these dyes provide evidence for redox processes and ROS formation, a potential drawback is that those probes act irreversibly by generating an accumulative signal rather than a reversible, dynamic response. Further, their lack of unambiguous subcellular localization and chemical specificity make it frequently difficult to draw mechanistic conclusions. During the last decade, genetically encoded biosensors have revolutionized the field of cell physiology by being targetable to specific subcellular compartments and enabling dynamic measurements (Morgan et al., 2016). Among them, Grx1-roGFP2 for sensing *E*_GSH_ (Marty et al., 2009) and roGFP2-Orp1 for sensing transient changes in H_2_O_2_ (Nietzel et al., 2019); the latter being based on a redox relay between the H_2_O_2_-sensitive Gpx3 peroxidase from yeast (Orp1) and roGFP2 (Gutscher et al., 2009). These sensors have become instrumental to monitor the dynamics of oxidative signals in real-time in a wide range of organisms, including plants. Similarly, probes of the HyPer family, which exploit the H_2_O_2_-sensitive bacterial transcription factor OxyR for their response, can report on local alterations in H_2_O_2_ concentrations (Bilan and Belousov, 2018; Pak et al., 2020).

Despite compelling evidence for the signaling functions of H_2_O_2_, it is neither known how H_2_O_2_ concentrations and the redox buffers dynamically respond to increased ROS production in chloroplasts nor how much other organelles contribute to a cumulative oxidation in the cytosol. Here, we targeted two different roGFP2-based biosensors to the stroma of the chloroplasts, the cytosol and the matrix of the mitochondria to live monitor the local *E*_GSH_ and H_2_O_2_ dynamics specifically in those three compartments. We investigated the dynamic subcellular responses to primary oxidative events triggered by MV, light or a combination of both. To dissect the contribution of chloroplasts and mitochondria in the MV-induced overall oxidation, the respective ETCs were blocked using ETC-specific inhibitors acting at early steps of electron transport.

## RESULTS

### Spectral properties of roGFP2-based probes *in planta*

To visualize changes of *E*_GSH_ or H_2_O_2_ levels in chloroplasts, cytosol and mitochondria, we selected previously published Arabidopsis reporter lines with roGFP2 linked to Grx1 or Orp1, respectively (Marty et al., 2009; Park et al., 2013; Albrecht et al., 2014; Nietzel et al., 2019). Since no roGFP2-Orp1 reporter line for H_2_O_2_ sensing in the plastid was available, we generated this line *de novo* (Supplemental Fig. S1). Subcellular localization of all reporter constructs was verified side-by-side in 7-day-old seedlings by confocal microscopy (Fig. 1A–F, left panels and Supplemental Fig. S1). Plants of the same age were used to systematically corroborate the *in vivo* excitation spectra of both redox sensors in all three compartments of intact seedlings. Sensor response and the dynamic spectroscopic response range was assessed by recording the fluorescence of seedlings immersed in imaging buffer using a fluorescence multiwell plate reader. Fluorescence spectra were collected for non-treated seedlings and seedlings incubated with either 20 mM DTT for complete reduction or with 100 mM H_2_O_2_ for complete oxidation of the sensors *in situ* (Fig. 1A–F, right panels). Sensor fluorescence intensities were sufficiently high to be clearly distinguishable from background fluorescence with a suitable signal-to-noise ratio for *in situ* readings (Supplemental Fig. S2). Fully reduced Grx1-roGFP2 (roGFP2-Grx1 in the case of the mitochondria) or roGFP2-Orp1 showed low excitation at 400 nm and a pronounced excitation peak close to 488 nm in all compartments. Probe oxidation led to the appearance of a second distinct excitation peak close to 400 nm, while excitation at 488 nm was decreased (Fig. 1A–F, right panels, Supplemental Fig. S2). The spectral behavior of both probes *in planta* was consistent with the spectra of the purified roGFP2 *in vitro* (Fig. 1H). These data validate that changes in the redox state of both roGFP2-based sensor variants can be reliably visualized and recorded in chloroplast stroma, cytosol and mitochondrial matrix using plate reader-based fluorimetry (Nietzel et al., 2019; Wagner et al., 2019).

**Figure 1.**
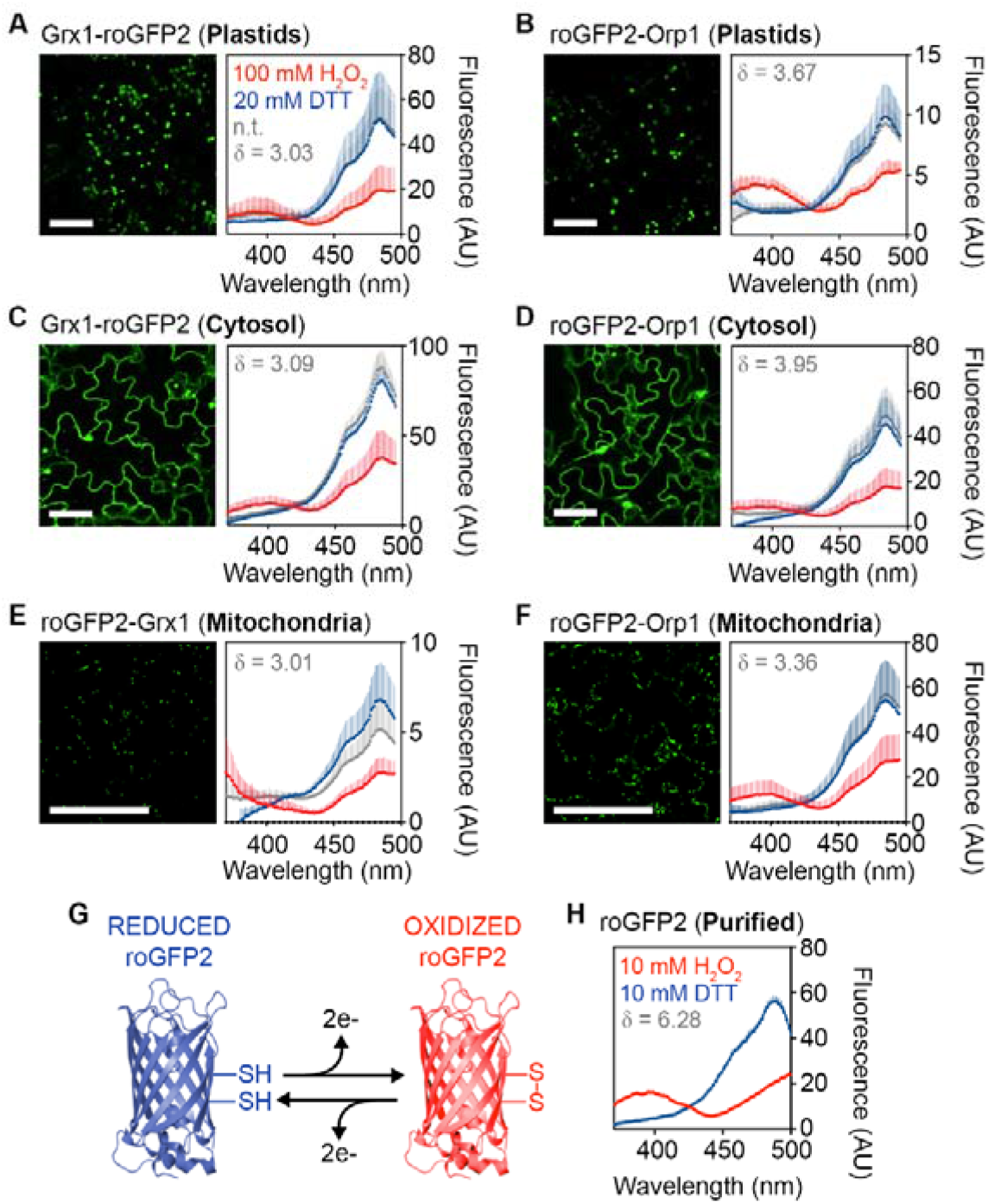
Subcellular localization and spectral behavior of *E*_GSH_ and H_2_O_2_ sensors in Arabidopsis. **A–F (left panels)**, Confocal microscopy images of leaf epidermal cells from 7-day-old seedlings stably expressing Grx1-roGFP2, roGFP2-Grx1 or roGFP2-Orp1 targeted to plastids (**A–B**), cytosol (**C–D**) or mitochondria (**E**–**F**). All images show roGFP2 fluorescence recorded with λ_ex_ = 488 nm and λ_em_ = 505-530 nm. Bars, 50 μm. **A–F (right panels),** Grx1-roGFP2, roGFP2-Grx1 or roGFP2-Orp1 fluorescence excitation spectra for non-treated seedlings (n.t., grey), and after reduction with 20 mM DTT (blue) or oxidation with 100 mM H_2_O_2_ (red). All spectra were recorded on a plate reader from 7-day-old seedlings with emission at 520±5 nm and using the same gain for all lines. The curves show the mean of the fluorescence in arbitrary units (AU) +SD, with *n* ≥ 3 biological replicates, where each replicate is an independent pool of 4–5 seedlings. All spectra were corrected for the autofluorescence measured in non-transformed control seedlings (see Supplemental Fig. S1). The dynamic range (δ) for the maximum change of the fluorescence ratio between the fully oxidized and fully reduced sensor was calculated from the fluorescence collected after sensor excitation at 410 and 480 nm. **G**, Schematic model of roGFP2 structure highlighting the disulfide bond formation upon reversible oxidation. **H**, Excitation spectrum of purified roGFP2 measured under similar conditions as the seedlings. To achieve full reduction and full oxidation, the purified protein was incubated in 10 mM DTT or 10 mM H_2_O_2_, respectively. Mean +SD, n = 6.

### Real-time monitoring of *E_GSH_* and H_2_O_2_ dynamics in Arabidopsis in response to externally imposed oxidative stress

To further validate the responsiveness of both probes *in planta*, 7-day-old Arabidopsis seedlings expressing cytosol-targeted Grx1-roGFP2 or roGFP2-Orp1 were exposed to different concentrations of H_2_O_2_ to impose oxidative stress (Fig. 2A, B). Changes in the redox state of both sensors were followed in real-time by exciting roGFP2 at 410 nm and 480 nm (Fig. 2B, C, E). The recorded fluorescence of roGFP2 in the individual channels (410 and 480 nm) showed an immediate response in opposite directions upon addition of H_2_O_2_, with a increase of the 410 nm channel while the fluorescence excited at 480 nm decreased (Fig. 2C, E; Supplemental Fig. S3A-E). After reaching a peak of oxidation, a gradual recovery of both channels towards the starting values occurred over a period of about four hours (Fig. 2C, E, Supplemental Fig. S3). For analysis, the 410 nm/480 nm fluorescence ratio was calculated and used as a measure for sensor oxidation (Fig. 2D, F). Starting from low ratio values in non-treated seedlings the addition of H_2_O_2_ caused a rapid increase of the 410 nm/480 nm fluorescence ratio. While the speed of oxidation was independent of the amount of H_2_O_2_, the maximum peak height increased with increasing concentrations of H_2_O_2_ (Fig. 2D, F). In all experiments, the minimum and maximum fluorescence ratios of the respective probes in their fully oxidized and fully reduced state were determined after each experiment as shown for the cytosolic roGFP2-Orp1 sensor (Supplemental Fig. S3F).

**Figure 2.**
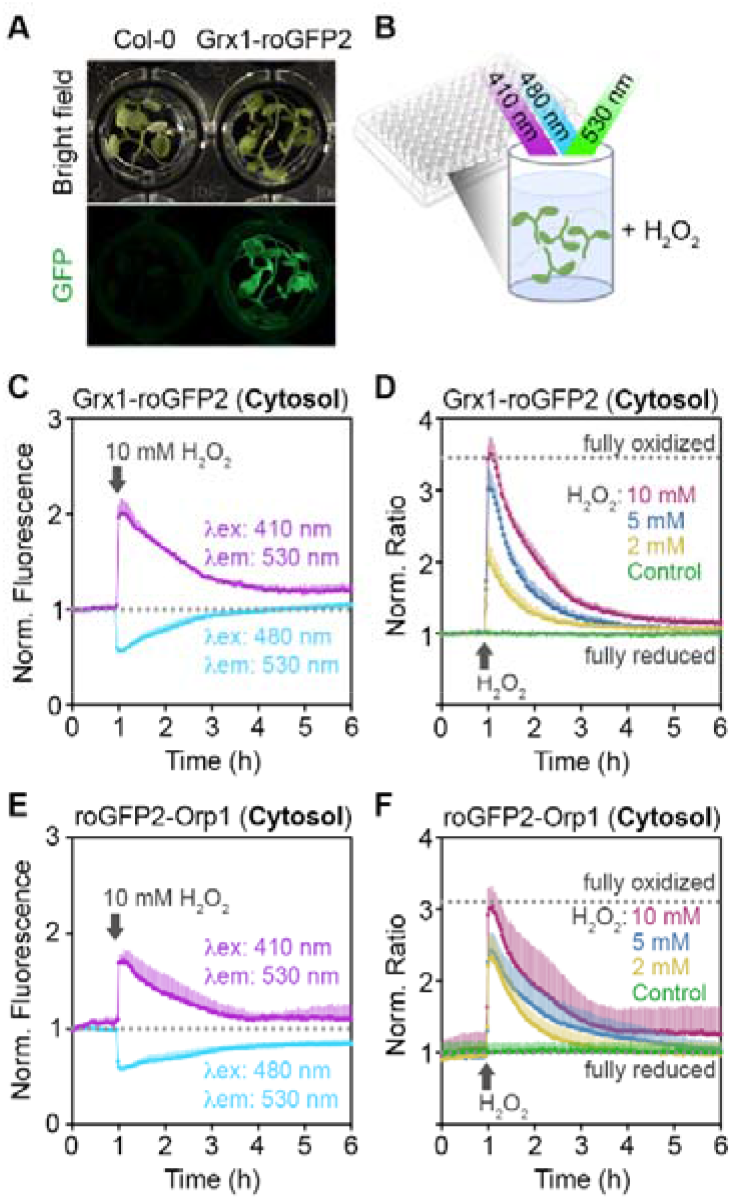
Real-time monitoring of cytosolic Grx1-roGFP2 and roGFP2-Orp1 redox changes upon imposition of oxidative stress *in planta*. **A,** Pools of 7-day-old seedlings (4–5 per well) expressing Grx1-roGFP2 or roGFP2-Orp1 in the cytosol were placed in a 96-well plate. **B,** The redox state of the sensors was measured as the roGFP2 fluorescence after sequential excitation at λ_ex_ = 410±5 nm and λ_ex_ = 480±5 nm. Fluorescence was always recorded at λ_em_ = 530±20 nm. **C, E,** Time–resolved fluorescence recordings for the two independent channels after addition of 100 mM H_2_O_2_ at t = 1 h. Both curves are normalized to their initial values before imposed oxidation (dotted lines). **D, F,** Fluorescence ratio values calculated to the original fluorescence in both channels in response to different concentrations of H_2_O_2_. For control samples only buffer was added. The curves show the mean ratio +SD from *n* = 4 biological replicates, where each replicate is an independent pool of 4–5 seedlings. The ratio values are normalized to the original ratio values before addition of H_2_O_2_. The experiment was repeated three times with similar results. Dotted lines in panels D and F indicate minimum and maximum ratio values measured from the same wells at the end of the experiment during incubation in 100 mM H_2_O_2_ for full oxidation and in 20 mM DTT for full reduction of the probes (see Supplemental Fig. S3).

### The dominant impact of MV-induced glutathione oxidation occurs in the chloroplasts

After confirming the fast and concentration-dependent response of the sensors towards H_2_O_2_ treatments *in vivo*, we evaluated the sensitivity of the plate reader-based fluorimetry setup to detect redox changes induced by MV. For this purpose, 7-day-old seedlings expressing Grx1-roGFP2, roGFP2-Grx1 or roGFP2-Orp1 targeted to plastid stroma, the cytosol or the mitochondrial matrix were treated with different concentrations of MV and sensor fluorescence was continuously recorded in plate reading mode for 6 h (Fig. 3). In chloroplasts, the oxidation of both probes gradually increased over time, with roGFP2-Orp1 showing a faster oxidation immediately after MV treatments compared to Grx1-roGFP2 (Fig. 3A–C; Supplemental Fig. S4A–B). In contrast to the response in chloroplasts, the fluorescence ratios of cytosolic Grx1-roGFP2 and roGFP2-Orp1 remained low suggesting that both probes remained largely reduced under the same treatments (Fig. 3D–F). Despite the slow oxidation, ratio values five hours after addition of MV were higher than in control seedlings. . In mitochondria both sensors revealed a gradual oxidation after the addition of 100 μM MV, roGFP2-Orp1 ratio increased more strongly during the first five hours after the addition of MV than roGFP2-Grx1(Fig. 3G–I). Because the action of MV in chloroplast is light-dependent, we further tested whether the flashes of light used for roGFP2 excitation were sufficient to cause the oxidation of roGFP2 by photooxidation effects. Increasing the cycle time of fluorescence readings from individual wells from 3 to 60 min abolished the detectable roGFP2-Orp1 oxidation in chloroplasts and the cytosol, which is consistent with light-dependency of MV-mediated ROS generation in chloroplasts (Supplemental Fig. S5). In mitochondria, the oxidation five hours after addition of MV was also visible albeit to a lower extent than with high flash frequencies. The steady increase of the fluorescence ratio with high repetition rates for the readout (Fig. 3) suggests that the oxidation effect caused by the herbicide is not reversible and exceeds the endogenous reducing capacity. The pronounced increase in the fluorescence ratio for both sensors targeted to chloroplast and mitochondria in response to 100 μM MV suggests that a local increase of H_2_O_2_ occurs in these organelles (Supplemental Fig. S4A–B), implying that the oxidation of the chloroplastic glutathione pool exceeds the oxidation measured in the cytosol and in mitochondria and that the light flashes used to excite the fluorophore enhance the MV-induced oxidation.

**Figure 3.**
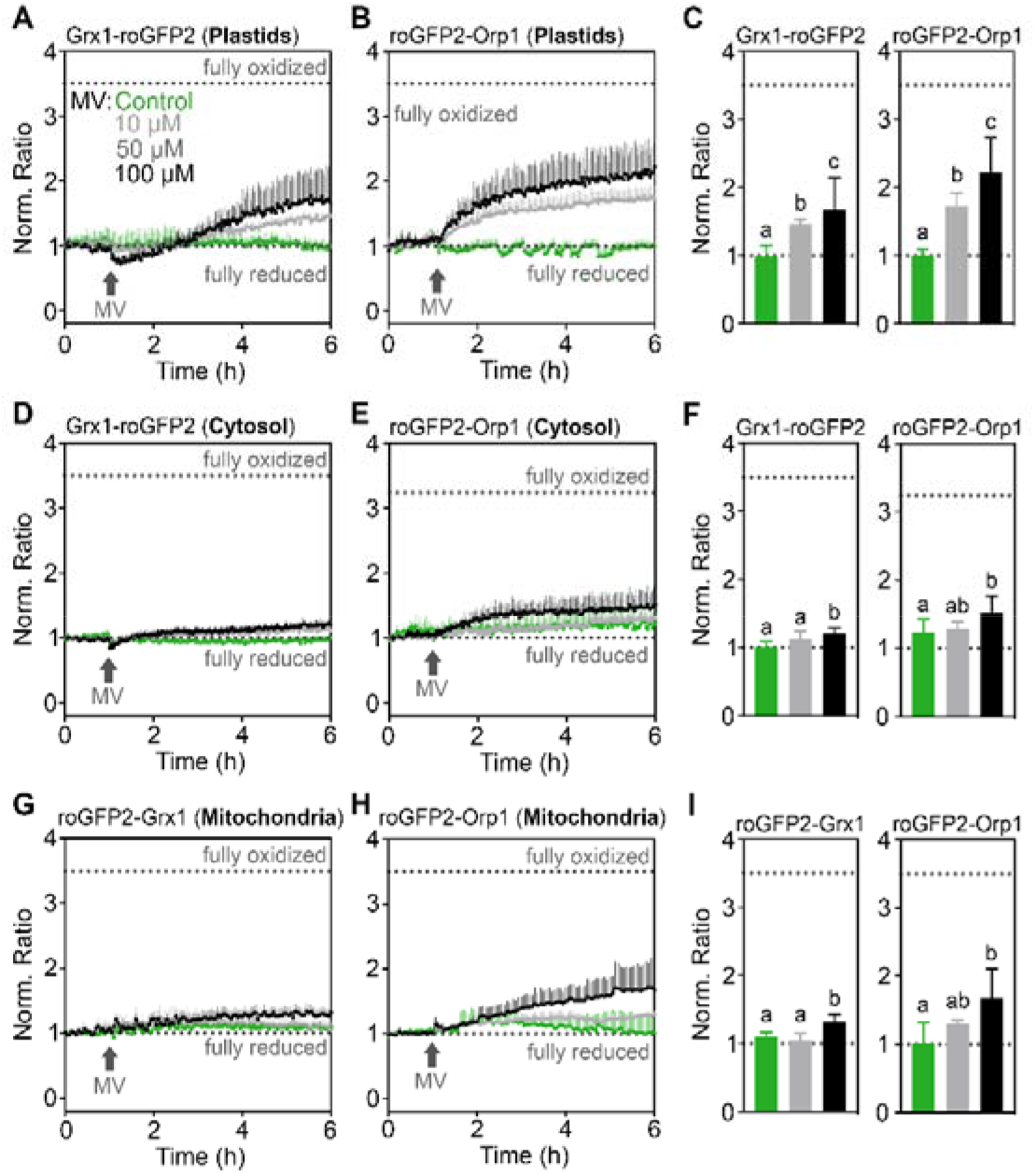
Real-time monitoring of *E*_GSH_ and H_2_O_2_ sensors upon MV-induced oxidation *in planta*. **A–I,** Seven-day-old seedlings stably expressing Grx1-roGFP2, roGFP-Grx1 or roGFP2-Orp1 targeted to the cytosol, plastids or mitochondria were placed in a 96-well plate with imaging buffer. After 1 h, MV was added to a final concentration indicated in panel A. In control samples (green), only buffer was added to maintain a uniform total buffer volume throughout the experiments. Ratio values were calculated from the fluorescence recorded by sequential excitation of probes at 410±5 nm and 480±5 nm, and normalized to the initial ratio at 0 h. Fluorescence was always recorded at 530±20 nm. Dotted lines indicate ratio values measured from the same wells at the end of each experiment after incubation in 20 mM DTT for full reduction or 100 mM H_2_O_2_ for full oxidation of the probes. **C**, **F**, **I**, Endpoint ratio values at 6 h extracted from panels A, B, D, E, G and H. Mean ratios +SD, *n* ≥ 3 biological replicates, where each replicate is an independent pool of 4–5 seedlings. Different letters indicate statistical differences between ratios after log10 transformation, according to one-way ANOVA with Tukey’s multiple comparison test (*P* < 0.05). Data for individual excitation channels are presented in Supplemental Fig. S4.

### Continuous light enhances MV-induced oxidation in chloroplasts, cytosol and mitochondria

The more pronounced oxidation induced by MV in chloroplasts compared to the cytosol and mitochondria (Fig. 3) was related to the unavoidable intermittent illumination during data collection (Supplemental Fig. S5). While this primary oxidation in chloroplasts was exploited for the following experiments, the results would only be physiologically meaningful as long as the seedlings do not get seriously damaged over the course of the experiment, especially when illuminated at high frequencies or even intermittent periods of continuous light. To test whether MV in combination with illumination had obvious effects on plant viability, Arabidopsis seedlings that had been exposed to excitation light every three minutes for a 15 hour fluorescence recording were taken out of the plates and transferred to agar plates for phenotype documentation. All seedlings that had been repeatedly illuminated with excitation light were still green and fully turgescent irrespective of the MV concentration (Supplemental Fig. S6A). Even if seedlings were illuminated with constant actinic light with an intensity of 200 μmol m^-2^ s^-1^ for 1 h after the first 2 hours of the 15 h time course, no obvious toxic effect of MV could be recognized macroscopically (Supplemental Fig. S6B). By contrast, seedlings kept under constant actinic light with an intensity of 200 μmol m^-2^ s^-1^ for 15 h showed loss of chlorophyll already with 10 μM MV and even more seriously with 100 μM MV, which caused complete bleaching (Supplemental Fig. S6C).

After confirming that 1 h illumination outside the plate reader does not severely damage the seedlings, we used this regime to further boost ROS formation in chloroplasts (Fig. 4A–I). The 1-hour illumination with constant actinic light with an intensity of 200 μmol m^-2^ s^-1^ caused a transient increase in the fluorescence ratio of both sensors targeted to chloroplasts and mitochondria in untreated control seedlings (Fig. 4A–B, G–H, green curves). For technical reasons, resuming roGFP measurements after intermittent illumination of seedlings outside the plate reader was only possible after a lag time of about 10 seconds post-illumination, since the plates needed to be transferred back into the reader. Conversely, the sensors in the cytosol in control plants did not respond to illumination alone (Fig. 4D–E, green lines). The combination of MV and illumination, however, induced an increase in the fluorescence ratio for both sensors in all three compartments, resulting in a transitory peak of oxidation lasting close to 10 minutes after illumination. The long-term measurement shows a gradual oxidation over time (Fig. 4). This oxidation was dependent on the concentration of MV applied, albeit with different amplitudes depending on the probe and on the compartment. While Grx1-roGFP2 in the cytosol showed a pronounced long-term ratio increase in plants pre-incubated with 50 and 100 μM MV, the ratio change of cytosolic roGFP2-Orp1 was limited to a minor reversible increase only (Fig. 4D–F). In mitochondria, light exposure caused transient oxidation of roGFP2-Orp1 and roGFP2-Grx1 albeit without pronounced differences between the two probes (Fig. 4G–I).

**Figure 4.**
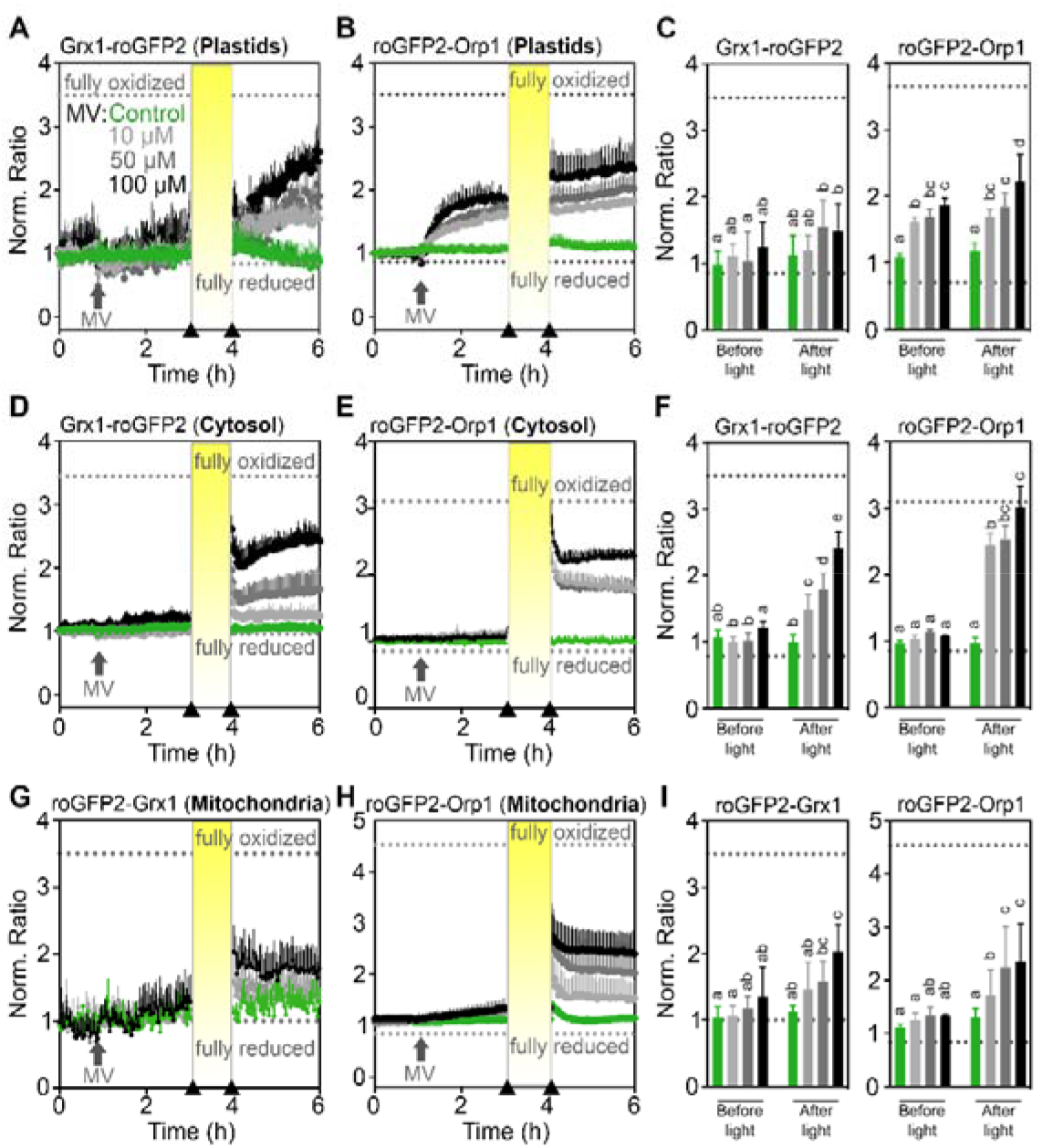
Light enhances MV-induced oxidation of roGFP2-derived redox sensors. **A–I,** Seven-day-old seedlings stably expressing the indicated sensor constructs in plastids, in the cytosol or in mitochondria were placed in a 96-well plate with imaging buffer. After 1 h, MV was added to final concentrations of indicated in panel A. Arrows on the x-axes of panels A, B, D, E, G, and H indicate the time points at which data for the bar charts in panels **C**, **F** and **I** were extracted (i.e. before and after illumination). In control samples (green), only buffer was added. Oxidation of the sensors was recorded as the normalized ratio of the fluorescence recorded with excitation at 410±5 nm and 480±5 nm, respectively. Fluorescence was always recorded at 530±20 nm. After a pre-incubation with MV for 2 h, seedlings were intermittently illuminated for 1 h with actinic light (200 μmol m^-2^ s^-1^) and redox measurements were subsequently resumed for 2 h. Dotted lines indicate ratio values measured from the same wells at the end of the experiment after incubation in 20 mM DTT for full reduction or 100 mM H_2_O_2_ for full oxidation of the probes. Mean ratios +SD, *n* ≥ 4 biological replicates, where each replicate is an independent pool of 4–5 seedlings. Different letters represent statistical differences between ratios after log10 transformation, according to one-way ANOVA with Tukey’s multiple comparison test (*P* < 0.05). Data for individual channels can be found in Supplemental Fig. S7.

### Chloroplasts and mitochondria contribute to MV-induced oxidative stress

Flash illumination for roGFP2 fluorescence measurements during plate reader-based fluorescence excitation with intervals of 1 h did not lead to changes in roGFP2 fluorescence ratios in chloroplasts and the cytosol (Supplemental Fig. S5B). In mitochondria, however, the fluorescence ratio did show a small increase pointing at the possibility of an autonomous mitochondrial oxidative response, which was also recently concluded by independent work using orthogonal approaches (Cui et al., 2019). To dissect the relative contributions of chloroplasts and mitochondria to the MV-induced oxidative response in the cytosol, we used inhibitors for the two different ETCs. Seedlings expressing the different sensors were pre-treated with either 3-(3,4-dichlorophenyl)-1,1-dimethylurea (DCMU) to inhibit the electron transport between PSII and plastoquinone, or rotenone to inhibit the mETC at complex I (Fig. 5A–B). DCMU pre-treatment decreased MV-induced oxidation of both plastid-targeted sensors compared to seedlings treated with MV alone (Fig. 5C, dark green and black curves). The inhibitory effect of DCMU was most evident immediately after illumination in seedlings expressing Grx1-roGFP2. In seedlings expressing roGFP2-Orp1, DCMU inhibited the oxidation of the sensor as well, albeit to a lesser extent. Inhibition of the pETC by DCMU also inhibited the light-induced oxidation of the cytosolic Grx1-roGFP2 (Fig. 5E) and mitochondrial roGFP2-Grx1 (Fig. 5G, left panel). At the same time the fluorescence ratio increase of mitochondrial roGFP2-Orp1 was restricted to about 50–70% by DCMU (Fig. 5G, right panel). Pre-treatments with rotenone noticeably suppressed the illumination- and MV-induced ratio increase of Grx1-roGFP2 in chloroplasts and cytosol (Fig. 5D, F, left panels, dark red and black curves). The inhibitory effect of rotenone, however, was less pronounced than that of DCMU (Fig. 5C, E, left panels).

**Figure 5.**
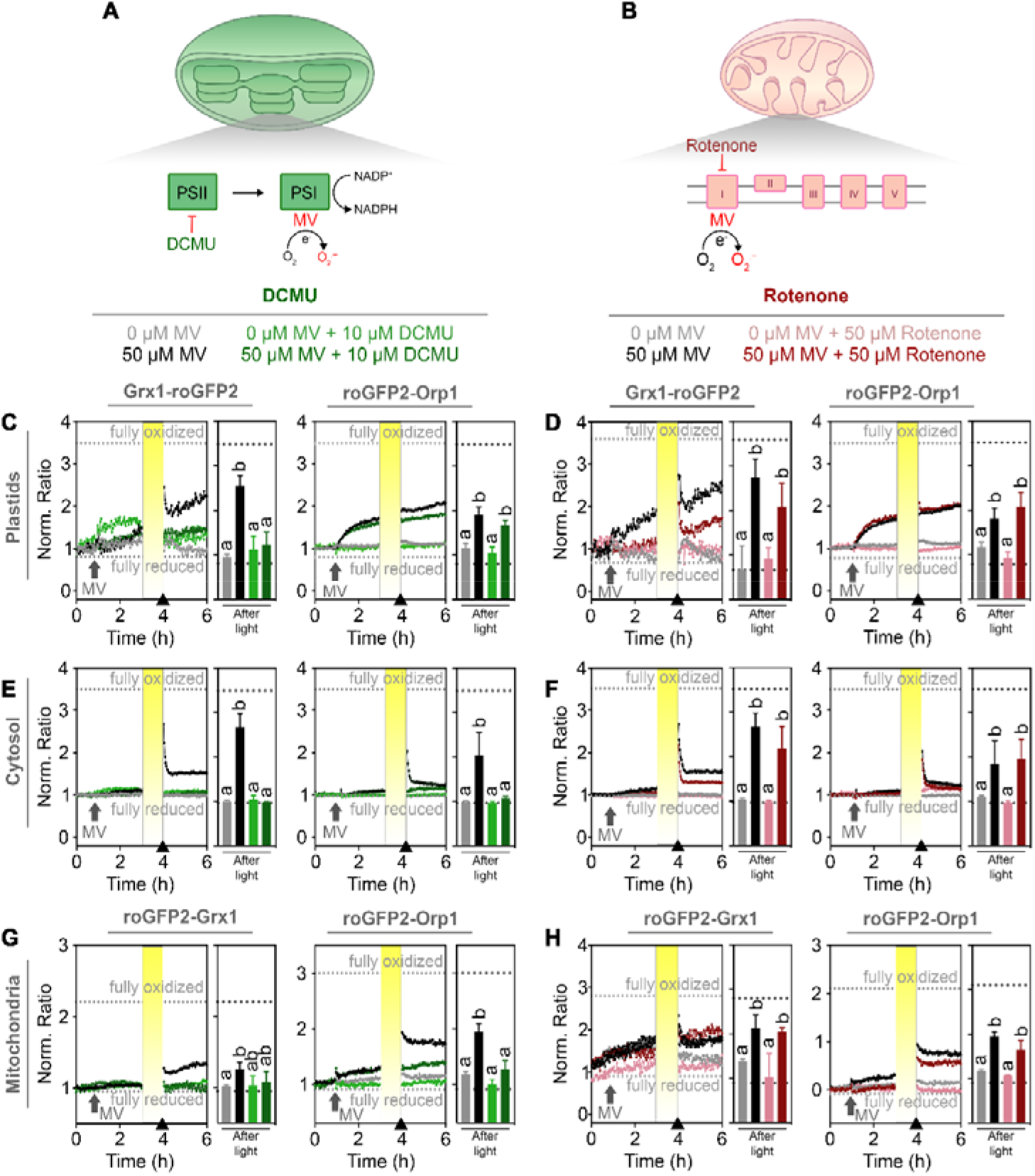
Contribution of chloroplast and mitochondrial ETCs to MV-induced oxidation. **A,** Model depicting the function and interplay of MV and ETC inhibitors on superoxide production in a chloroplast. MV withdraws electrons (e^-^) from photosystem I (PSI) and transfers them to molecular oxygen (O_2_) to form superoxide (O_2_^·-^). DCMU is an inhibitor that specifically blocks electron transfer from photosystem II (PSII) to plastoquinone. **B,** In the mitochondrion, MV is able to transfer electrons from complex I of the mitochondrial electron transport chain to O_2_, generating O_2_^·-^. Rotenone inhibits complex I activity. **C–H,** Seven-day-old seedlings stably expressing the indicated sensor constructs in plastids, cytosol or mitochondria were placed in 96-well plates with imaging buffer as a control or buffer supplemented with 10 μM DCMU or 50 μM rotenone to inhibit the electron flux along the chloroplastic or mitochondrial ETCs, respectively. After 1 h, MV was added to a final concentration of 50 μM or buffer as a control. After 2 h of treatments, the samples were intermittently exposed to 1 h of actinic light (200 μmol m^-2^ s^-1^). Data indicate the mean normalized ratio of the sensor fluorescence sequentially excited at 410±5 nm and 480±5 nm, and collected at 530±20 nm in at least four biological replicates (left panels). Arrows on the x-axes of panels C–H indicate the time point at which data for the bar charts (right panels). Dotted lines indicate ratio values measured from the same wells at the end of the experiment during incubation in 20 mM DTT for full reduction or 100 mM H_2_O_2_ for full oxidation of the probes. Different letters represent statistical differences between ratios after log10 transformation, according to one-way ANOVA with Tukey’s multiple comparison test (P < 0.05).

By contrast, roGFP2-Orp1 targeted to the plastids and the cytosol were not affected by incubation with rotenone (Fig. 5D, F, right panels). Rotenone pre-treatment on plants expressing the two sensors in mitochondria showed minor inhibition of the MV-induced ratio increase after illumination, particularly for the fluorescence ratio of roGFP2-Orp1 (Fig. 5H).

Immediately upon addition of MV to seedlings, a slight oxidation of the sensor in both chloroplasts and mitochondria occurred independently of the illumination treatment (Fig. 5C, D, G, H). The oxidation was still visible in chloroplasts and mitochondria of seedlings pre-treated with DCMU, but was less pronounced in seedlings pre-treated with rotenone (Fig. 5H). This confirms that the pETC largely contributes to the light-induced oxidation of all three compartments, while the mETC contributes only to a minor extent after illumination.

## DISCUSSION

### Dynamic recording of oxidative processes in multiple subcellular compartments of plant tissues using plate reader-based fluorimetry

Redox-sensitive GFPs have paved the way to elucidate the distinct differences in local *E*_GSH_ of subcellular compartments in plant cells (Jiang et al., 2006; Meyer et al., 2007; Schwarzländer et al., 2008). While roGFP2-based sensors for *E*_GSH_ are highly robust and reliable, sensor constructs for H_2_O_2_ monitoring are more diverse and exhibit a combination of integrated features (Schwarzländer et al., 2016). Probe variants of the HyPer family have been used multiple times in different subcellular compartments (Costa et al., 2010; Boisson-Dernier et al., 2013; Exposito-Rodriguez et al., 2017; Rodrigues et al., 2017; Mangano et al., 2018). Because HyPer is based on a circular permuted YFP a cleft in the artificial barrel structure allows direct access of protons to the central chromophore rendering most HyPer probes highly sensitive to pH, a feature that could potentially cause major artefacts in light-dependent measurements in chloroplasts (Belousov et al., 2006; Schwarzländer et al., 2014). An updated version, HyPer7, has been proven to be pH insensitive, but has not yet been used in plant research (Pak et al., 2020). Fusions of roGFP2 with peroxidases like the glutathione peroxidase Orp1 or peroxiredoxins, such as Tsa2 overcome this limitation because the roGFP2 ratio is pH-insensitive over the whole physiological range (Schwarzländer et al., 2008; Gutscher et al., 2009; Morgan et al., 2016). Orp1 fused to roGFP2 was chosen in this work based on its sensitivity to low H_2_O_2_ levels (Delaunay et al., 2002; Sobotta et al., 2013), its pH-insensitivity, and its dependence on reduction via Grx and GSH (Nietzel et al., 2019).

Environmental stress conditions can trigger oxidative processes via ROS formation. However, oxidation occurs with different dynamics and amplitudes in different subcellular compartments (Schwarzländer et al., 2009; Rosenwasser et al., 2011; Bratt et al., 2016). Dynamic measurements after triggering oxidative stress have been successfully carried out in the cytosol and mitochondria of cells and tissues placed in perfusion chambers on fluorescence microscopes (Schwarzländer et al., 2009). While high throughput approaches are limited in a typical microscopy set-up, plate reader-based approaches have enabled dynamic long-term measurements with multiple parallel samples in one experiment (Rosenwasser et al., 2010; Rosenwasser et al., 2011; Bratt et al., 2016; Nietzel et al., 2019; Wagner et al., 2019). Comparative measurements with Grx1-fused roGFP2 for *E*_GSH_ and roGFP2-Orp1 for H_2_O_2_ sensing recently revealed differential responses of both probes in the cytosol and the mitochondrial matrix (Nietzel et al., 2019). Our results show that this approach can be extended to the chloroplasts, as the predominant generators of ROS under illumination (Mubarakshina et al., 2010).

While genetically encoded biosensors can be powerful tools for *in vivo* monitoring of physiological parameters, potential problems caused by overexpression, mis-targeting or incomplete targeting have been observed (Albrecht et al., 2014; De Col et al., 2017). In contrast to a slight developmental delay that we observed previously for Arabidopsis plants with mitochondrial roGFP2-Orp1 (Nietzel et al., 2019), Grx1-roGFP2 and roGFP2-Orp1 can both be targeted to plastids without any mis-targeting and without causing any apparent developmental phenotype. The absence of obvious phenotypes suggests that the reporter constructs and their import do not interfere with normal plastid functions. In microscopic experiments with high spatial resolution incomplete targeting may not pose a major problem or can even be exploited as an advantageous feature for simultaneous recording of physiological responses in two compartments (Marty et al., 2019). As we employ plate reader assays in which only the overall fluorescence from a biological sample is recorded, correct targeting of the probes to all compartments is of utmost importance and was carefully validated.

Previous reports using roGFP-based sensors have demonstrated a fast and reversible oxidation in response to external H_2_O_2_ (Meyer et al., 2007; Morgan et al., 2011). We observed similar fast oxidation kinetics for both tested sensors in the cytosol of Arabidopsis seedlings. 10 mM H_2_O_2_ were sufficient to reach the maximum oxidation of the sensors. This maximum oxidation, however, was not maintained but rather followed by an immediate recovery towards the fully reduced state over the course of five hours. This decrease in the fluorescence ratios of roGFP2 after severe oxidative challenge indicates a decrease of intracellular H_2_O_2_ and re-reduction of the cytosolic glutathione buffer, respectively. This recovery shows the remarkable efficiency of the plant peroxide detoxification machinery, enabling 4-5 seedlings with a total fresh weight of ~15 mg to clear a total volume of 200 μL from 10 mM H_2_O_2_ within five hours. This would amount to an average detoxification of about 440 nmol (g FW)^-1^ min^-1^, which interestingly is in the order of 50-5,000 nmol (g FW)^-1^ that has been reported for the H_2_O_2_ content in unstressed leaves (Queval et al., 2008). Although it is not the focus of this study, our experimental setup thus appears well suited for further genetic dissection of cellular peroxide detoxification systems (Smirnoff and Arnaud, 2019). The transitory peak for the maximum oxidation also highlights the need for immediate measurement of complete sensor oxidation for calibration purposes (Schwarzländer et al., 2008).

### Real-time monitoring of MV-induced oxidative stress

While both roGFP2-based probes used here show an oxidative response after MV treatment within hours, it is already known that exposure of plants to MV triggers distinct changes in gene expression. In cucumber, MV causes the accumulation of ROS and lipid peroxides after 1 h, while GSH oxidation and an increase in ascorbate peroxidase (APX) and glutathione peroxidase (GPX) activities were identified 48 h after MV application (Liu et al., 2009). Light acts as an enhancer of the MV-induced oxidation, but is not essential for MV-induced oxidative damage (Cui et al., 2019; Shapiguzov et al., 2019).

Plants frequently contain a broad range of autofluorescent endogenous compounds (Müller et al., 2013). Reliable interpretation of sensor responses thus depends on correct recording and subtraction of such signals underlying and potentially obscuring the true roGFP2 signal (Fricker, 2016). While for short-term treatments with oxidative changes induced by light and/or MV this can be done reliably and additionally controlled for by careful monitoring the raw data for each individual channel, deviations in long-term measurements cannot be fully excluded. For long-term recordings we found that in some cases MV treatment led to strong changes of the apparent ratio recorded for both roGFP2-based probes *in vivo* even though the individual channels showed little change. Although we cannot fully explain the exact kinetics of the fluorescence ratio recorded over several hours after MV treatment, we show that the reported effects of MV and light are reliable and reproducible. In addition, inhibitor treatments confirmed that the observed effects are causally connected to primary oxidation occurring in the respective subcellular compartments.

Treatment of seedlings with MV without additional illumination caused oxidation in chloroplasts, and to a minor extent in mitochondria and the cytosol. This contrasts with earlier measurements in which MV caused a minor oxidation in mitochondria and a pronounced oxidation in the cytosol of Arabidopsis cotyledons (Schwarzländer et al., 2009). A major difference between the two experiments is the use of a confocal microscope with laser excitation targeting a small number of cells by Schwarzländer et al. (2009), while a plate reader with less intense excitation light collecting fluorescence from whole seedlings was used in this work. In the first 2 h after MV addition, the response of the roGFP2-Orp1 sensor was faster and more pronounced compared to Grx1-roGFP2 in plastids or roGFP2-Grx1 in mitochondria, which indicates an increase of H_2_O_2_ before a change in *E*_GSH_.However, because the oxidized roGFP2-Orp1 depends on GSH for its reduction and because the roGFP2 domain may on its own react with GSH/GSSG, a gradual change in sensor oxidation may also reflect changes in the *E*_GSH_. If both sensors react to imposed stress (i.e. 5 h after MV addition) it is thus not possible to dissect whether the observed oxidation is caused by H_2_O_2_ production or an increase of *E*_GSH_ (Meyer and Dick, 2010; Nietzel et al., 2019). We observed complete oxidation of plastid-targeted roGFP2-Orp1 and partial oxidation of plastid-targeted Grx1-roGFP2 after MV addition, indicating that the excitation light in the plate reader is still sufficient to trigger electron flux in the pETC and hence the formation of ROS (Supplemental Fig. S5). The MV-induced oxidation was considerably slower than the oxidation induced by incubation of seedlings in H_2_O_2_ (Fig. 3), in accordance with only intermittent illumination during plate reader measurements. The minor oxidation observed for cytosolic sensors after MV challenge indicates that either low amounts of ROS are leaving the chloroplasts under these conditions, or that the capacity of the cytosolic scavenging systems is sufficient to detoxify the H_2_O_2_ leaking from chloroplasts. H_2_O_2_ produced in chloroplasts may be detoxified locally through the glutathione–ascorbate cycle. The *E*_GSH_ in each compartment is independent and not directly correlated (Marty et al., 2019). The resultant GSSG is contained within the organelles, leading to a local change in *E*_GSH_ to less reducing values. The lethal phenotypes of Arabidopsis mutants deficient in plastidic GR2 strongly suggest that GSSG cannot be efficiently exported from plastids (Marty et al., 2019). In addition, stromal *E*_GSH_ responds dynamically to light (Haber and Rosenwasser, 2020; Müller-Schüssele et al., 2020) and GSSG can be formed via several enzymes directly or indirectly involved in ROS scavenging (DHAR, GRX, PrxII, MSRB, among others), and its efficiently recycled by GR2.

### ROS as a putative mobile signal between cellular sub-compartments

In photosynthetic organisms, light exposure triggers activation or inactivation of multiple redox-regulated enzymes containing thiol-switches (Cejudo et al., 2019). In addition, 20–60 seconds of light stress in Arabidopsis leads to an increase in GSH and total glutathione through processes associated with high levels of nitric oxide and photorespiratory processes providing increased amounts of GSH precursors (Choudhury et al., 2018). From high light treatments of plants transiently expressing HyPer it was deduced that oxidation in plastids and the nucleus takes less than one second (Exposito-Rodriguez et al., 2017). In our experiments, we show a change in the oxidation of sensors targeted to plastids and mitochondria in plants exposed to light (Fig. 4A–B G–H, green curves), indicating that light exposition changes the redox homeostasis in these compartments. Further research will be needed to increase the time resolution of these light-induced redox changes, especially using pH-insensitive sensors such as roGFP2-Orp1 and Grx1-roGFP2, since pH adjustments may be challenging for fast events.

Light exposure of samples pre-treated with MV increased the oxidation in chloroplasts, cytosol and mitochondria more than MV alone (Fig. 4), in accordance with enhanced MV toxicity due to pETC activation (Cui et al., 2019). Oxidation of the Grx1-roGFP2 and roGFP2-Grx1 in all compartments was dependent on the pETC, as it was inhibited by DCMU. This was the same for roGFP-Orp1 targeted to cytosol and mitochondria, while only a minor inhibition was observed in plastids (Fig. 5C, E, G). This may indicate that the concentration of DCMU used still allows H_2_O_2_ production in the chloroplast or that it enhances the production of peroxides through production of singlet oxygen (Fufezan et al., 2002; Krieger-Liszkay, 2005; Dietz et al., 2016). In this same sensor line, no oxidation peak was observed immediately after light exposure (Fig. 4). This peak might have been missed due to a slightly slower transfer of the well plate back into the plate reader.

The oxidation of roGFP2-Orp1 in mitochondria after light and MV treatment was dependent on pETC activity, as it was inhibited by DCMU (Fig. 4G, right panel). This indicates that ROS produced in the chloroplast is escaping the local scavenging system and affects redox homeostasis in the cytosol and mitochondria. In high-light stress conditions, the association between plastids and nucleus increases (Exposito-Rodriguez et al., 2017), potentially fostering a direct transfer of ROS from the plastids to the nucleus. Chloroplastic H_2_O_2_ is believed to act as a secondary messenger involved in retrograde signalling to the nucleus (Chan et al., 2016), where it can modulate the transcriptome (Sewelam et al., 2014). As an example, the over-excitation of PSI can lead to ROS-induced changes in the redox state of antioxidant-related transcription factors such as ZAT10 (Rossel et al., 2007) and HSFA1D, the latter can enhance the expression of the peroxidase AXP2 (Jung et al., 2013).

The transfer of ROS from plastids to the cytosol might be facilitated by aquaporins (Mubarakshina et al., 2010). In mammals, Aqp8 and Aqp11 have been already characterized as essential ROS transporters from mitochondria and the endoplasmic reticulum, respectively (Chauvigné et al., 2015; Bestetti et al., 2020). In tobacco chloroplasts, the aquaporin NtAQP1 has been described as a gas pore for CO_2_ (Uehlein et al., 2008), while plant mitochondria harbour at least 1 aquaporin, TTIP5;1, which is still uncharacterized (Bienert and Chaumont, 2014). Dynamic recording of ROS formed at the chloroplastic and mitochondrial ETCs and downstream oxidative changes in other subcellular compartments as established in this work presents opportunities for future research regarding the role of aquaporins in facilitating ROS diffusion across membranes.

In mammals and other non-photosynthetic organisms, mitochondria constitute the main source of ROS production under MV treatments (Cochemé and Murphy, 2008). Our findings showed mild oxidation in the mitochondrial matrix after addition of MV (Fig. 3G–I), which can also be observed in the fluorescence intensity shifts of both individual channels at the moment of MV addition to seedlings expressing the mitochondria-targeted roGFP2-Orp1 (Supplemental figure S4F, supplemental figure S5F). This initial oxidative shift even before the light treatment was inhibited in plants pre-treated with the mETC inhibitor rotenone but not with the pETC inhibitor DCMU (Fig. 5H, right panels). This indicates that MV-toxicity in mitochondria is independent from the chloroplastic pETC. Notably, rotenone also inhibited the response of *E*_GSH_ to light/MV treatment in plastids and the cytosol (Fig. 5D, F, left panels). In isolated barley thylakoids rotenone inhibits a NAD(P)H dehydrogenase-like enzyme (NDH) which is implicated in cyclic electron transport via photosynthetic complex I (Teicher and Scheller, 1998). Despite its original name, NDH has been recently shown to preferentially accept electrons from ferredoxin rather than NADPH (Schuller et al., 2019). The key role of NDH in cyclic electron transport may nevertheless explain a direct modulation of light-induced MV toxicity by rotenone. In addition, mitochondria and chloroplasts are metabolically coupled (Schöttler and Tóth, 2014; Shameer et al., 2019). Therefore, we cannot discard an indirect inhibition of the pETC by decreasing the mETC with rotenone.

Efficient inhibition of the mETC with rotenone and the pETC with DCMU led to concomitant abolishment of MV-induced changes in both sensors. This observation further supports the notion that the responses of both probes are causally connected to the activity of either ETCs. The interpretation of alterations in roGFP2 fluorescence as a measure of oxidative responses linked to deviations of electrons from the ETCs thus appears valid (Fig. 6). Similarly, inhibitor-dependent abolishment of oxidation in the cytosol caused by MV in conjunction with light strongly suggests that the observed cytosolic alterations mirror the release of ROS from chloroplasts into the cytosol and mitochondria. With the respective probes and the plate reader setup it is thus possible to measure the immediate and dynamic oxidative response to a stress imposed on chloroplast, in the cytosol and mitochondria. Pulsed short-term stresses like e.g. short periods of strong illumination are followed by a recovery phase. Only if the stress occurs at higher frequencies or even persist permanently, this results in a long-term oxidation.

**Figure 6.**
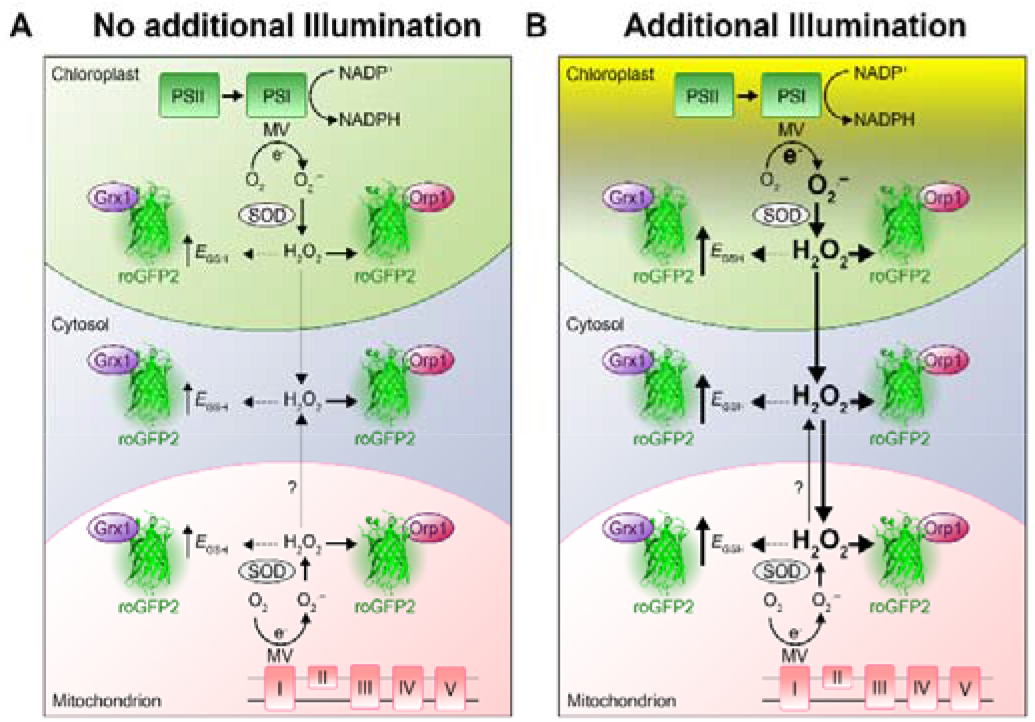
Increased chloroplast-derived ROS production caused by MV modulates the glutathione redox potential in chloroplasts, cytosol and mitochondria. **A,** MV is an herbicide that causes photo-oxidative stress in chloroplasts by diverting electrons from the photosystem I (PSI) to molecular oxygen (O_2_), leading to formation of superoxide (O_2_^·-^). In addition, MV is able to transfer electrons from complex I of the mitochondrial electron transport chain to O_2_, also generating O_2_^·-^. Superoxide is dismutated to H_2_O_2_ by chloroplastic or mitochondrial superoxide dismutases (SODs). Increased accumulation of H_2_O_2_ can lead to oxidation of the glutathione buffer, most likely via detoxification along the glutathione–ascorbate cycle (not shown). **B,** Additional light enhances electron transport in the chloroplasts, inducing the production of O_2_^·-^ and in consequence also H_2_O_2_. If the scavenging capacities of the chloroplast are surpassed, H_2_O_2_ might be leaking to the cytosol and mitochondria. H_2_O_2_ can indirectly increase local *E*_GSH_, which can be tracked using Grx1-roGFP2. The increase of H_2_O_2_ can be tracked using the oxidation of the roGFP2-Orp1 sensor.

### Concluding remarks

In this work, we have established a simple semi high-throughput approach to follow the contribution of different subcellular compartments to a ROS-mediated oxidation response in living tissues for multiple seedlings and treatment regimes in parallel. The results identify chloroplasts as the principal source of ROS in response to MV in illuminated, photosynthetically active tissue but also highlight the contribution of mitochondria to MV toxicity within plant cells. The ability to record dynamic redox-related changes will pave the way to better understand the interplay between redox imbalances in distinct plant compartments. In future experiments, measuring coupled redox dynamics in combination with genetics will enable further dissection of the signalling events between subcellular compartments.

## MATERIALS AND METHODS

### Plant material and growth conditions

*Arabidopsis thaliana* Col-0 ([L.] Heynh.) plants were obtained from NASC (www.arabidopsis.info). Transgenic Col-0 lines expressing the Grx1-roGFP2 sensor in the cytosol have been described earlier by (Marty et al., 2009). For targeting Grx1-roGFP2 to plastids, the sensor construct was cloned behind the target peptide of transketolase (TK_TP_) (Wirtz and Hell, 2003; Schwarzländer et al., 2008; Speiser et al., 2018). For measurements of *E*_GSH_ in the mitochondrial matrix, Col-0 plants were transformed with roGFP2-Grx1 cloned behind the target peptide of serine hydroxymethyltransferase (SHMT_TP_) as reported in (Albrecht et al., 2014) and (Marty et al., 2019). Cytosolic and mitochondrial targeted versions of the roGFP2-Orp1 sensor were previously reported (Nietzel et al., 2019), while the plant line expressing the plastid localized version of the sensor was generated in this work. For experiments with whole seedlings, seeds were surface-sterilized with 70% (v/v) ethanol, rinsed 3 times with sterile deionized water and stratified for 48 h at 4°C. Seeds were then sown on plates with 0.5x Murashige and Skoog growth medium (Murashige and Skoog, 1962) (Duchefa Biochemie, Haarlem, The Netherlands) supplemented with 0.1% (w/v) sucrose, 0.05% (w/v) MES (pH 5.8, KOH) and 0.8% (w/v) agar. Plates were incubated vertically in a growth chamber under a long-day regime (16 h light, 22 ± 2°C; 8 h dark 18 ± 2°C) with a photon flux density of 100 μmol m^-2^ s^-1^ for 7 days.

### Cloning of plastid-targeted roGFP2-Orp1 sensor and generation of transgenic plant lines

The roGFP2-Orp1 sequence was amplified by PCR from pBSSK:roGFP2-Orp1 (Gutscher et al., 2009) and fused to TK_TP_(as described in (Nietzel et al., 2019). Primers for roGFP2-Orp1 amplification were AACCATAGAGAAAACTGAGACTGCGGTGAGCAAGGGCGAGGAGCTGTTC and GTACAAGAAAGCTGGGTTCTATTCCACCTCTTTCAAAAGTTCTTC and for amplification of the targeting peptide TACAAAAAAGCAGGCTTCACCATGGCGTCTTCTTCTTCTCTCACT and GAACAGCTCCTCGCCCTTGCTCACCGCAGTCTCAGTTTTCTCTATGGTT. Fusion of both constructs was achieved by amplification with the primers GGGGACAAGTTTGTACAAAAAAGCAGGCTTCACC and GGGGACCACTTTGTACAAGAAAGCTGGGTTCTA. For constitutive plant expression (CaMV 35S promoter), the amplicon was cloned into pDONR207 (Invitrogen Ltd, Carlsbad, CA) and then into pH2GW7 (Karimi et al., 2002) using Gateway cloning (Invitrogen Ltd, Carlsbad, CA).

### Confirmation of subcellular roGFP2 targeting by confocal microscopy

Seven-day-old seedlings were imaged using a confocal laser scanning microscope (Zeiss LSM 780, connected to an Axio Observer.Z1; Carl Zeiss Microscopy, Jena, Germany) with a 40x lens (C-Apochromat 40x/1.2 W Korr). GFP and chlorophyll fluorescence were measured by excitation at 488 nm and emission at 505–530 nm (GFP) and 650–695 nm (chlorophyll). For mitochondrial counter staining, seedlings were vacuum infiltrated for 30 min on 200 nM MitoTracker Orange (Thermo Fisher Scientific, Waltham, MA) and measured by excitation at 543 nm and emission at 570-623 nm.

### Purification of recombinant roGFP2 variants

*E. coli* HMS174/Origami cells containing the pET30-roGFP2-His vector were cultured in liquid LB medium supplemented with 50 μg mL kanamycin at 37°C to an OD_600_ of 0.6–0.8. roGFP2-His expression and cell lysis were performed as described in (Nietzel et al., 2019). The lysate was centrifuged at 19,000 *g* for 15 min at 4°C and the supernatant filtered through a sterile filter of 0.45 μm nominal pore size. The filtered fraction was then loaded onto a Ni-NTA HisTrapTM column (GE Healthcare, Little Chalfont, UK) using a peristaltic pump at a flow rate of 1 mL min. Proteins were eluted from the column with a 10–200 mM imidazole gradient (100 mM Tris-HCl, pH 8.0, 200 mM NaCl) using an ÄKTA Prime Plus chromatography system (GE Healthcare). Fractions were collected and stored at 4°C.

### Spectral measurement of roGFP2 probe variants *in planta*

4–5 seedlings were placed in 250 μL imaging buffer (10 mM MES, 10 mM MgCl2, 10 mM CaCl2, 5 mM KCl, pH 5.8) in transparent Nunc^®^ 96-well plates. Samples were excited at 370–496 nm with a step width of 1 nm and the emission collected at 530±5 nm using a CLARIOstar plate reader (BMG Labtech, Offenburg, Germany). To achieve complete reduction or oxidation of the sensor, the imaging buffer was supplemented with either 20 mM 1,4-dithiothreitol (DTT) or 100 mM H_2_O_2_. Non-transformed wild-type seedlings were treated in the same conditions and used to determine the autofluorescence that was subtracted from the fluorescence recorded in roGFP2 lines. The spectral properties of recombinant roGFP2 were measured under the same conditions.

### Time-resolved ratiometric analysis of probe fluorescence *in planta*

All *in planta* measurements for all different probes were conducted in either a CLARIOstar or POLARstar plate reader (BMG Labtech). roGFP2 was excited by a filter-based excitation system at 410±5 nm and 480±5 nm. Fluorescence was collected using either a 530±20 filter for the POLARstar, or a 520±5 nm for the CLARIOstar. The fluorescence ratio was calculated as 410 nm / 480 nm and normalized to the ratio value at 0 h. Seedlings were treated with different concentrations of H_2_O_2_ in a final volume of 250 μL imaging buffer. To induce oxidative stress in seedlings by endogenous ROS production, MV (Sigma-Aldrich, Steinheim, Germany) was added at different final concentrations using the built-in automated injectors. Actinic light treatments were performed by placing the plate with seedlings pre-exposed to MV for 2 h under white LEDs with a photon flux density of 200 μmol m^-2^ s^-1^ for 1 h. Subsequently, the recording of roGFP2 fluorescence was continued in the dark (except for the short light flashes required for the measurements) within the plate reader. To assess the dynamic range of the probes *in planta* and for sensor calibration, 20 mM DTT and subsequently 100 mM H_2_O_2_ were added at the end of each experiment to fully reduce and fully oxidize the sensors. Between these treatments, samples were rinsed twice with imaging buffer. In each experiment, at least 4 technical replicates consisting of 4–5 pooled seedlings per well were used. Each experiment was repeated at least 3 times.

Samples were incubated with inhibitors of chloroplast and mitochondrial ETC prior to the addition of MV. To inhibit photosystem II (PSII), samples were treated with 10 μM 3-(3,4-dichlorophenyl)-1,1-dimethylurea (DCMU) dissolved in ethanol, whereas 50 μM rotenone (Sigma-Aldrich) was dissolved in dimethyl sulfoxide (DMSO) to inhibit complex I of the electron transport chain in mitochondria.

## SUPPLEMENTAL MATERIAL

**Supplemental Figure S1**. Subcellular localization of roGFP2-based probes for *E*_GSH_ and H_2_O_2_ in Arabidopsis.

**Supplemental Figure S2.** Raw fluorescence of Arabidopsis *E*_GSH_ and H_2_O_2_ sensors lines compared with non-transformed Col-0 plants.

**Supplemental Figure S3.** Calibration procedure to determine the dynamic range of the sensor exemplified for cytosolic roGFP2-Orp1.

**Supplemental Figure S4.** Individual excitation channels of roGFP2 fluorescence upon MV-induced oxidation in planta.

**Supplemental Figure S5.** Effect of the excitation light on the MV-induced oxidation of the roGFP2-Orp1 sensor.

**Supplemental Figure S6.** Effect of extended treatment of Arabidopsis seedlings with MV.

**Supplemental Figure S7.** Individual excitation channels of roGFP2 fluorescence upon light-enhanced MV-induced oxidation in planta.

